# Small females prefer small males: size assortative mating in *Aedes aegypti* mosquitoes

**DOI:** 10.1101/328930

**Authors:** Ashley G. Callahan, Perran A. Ross, Ary A. Hoffmann

## Abstract

With *Aedes aegypti* mosquitoes now being released in field programs aimed at disease suppression, there is interest in identifying factors influencing the mating and invasion success of released mosquitoes. One factor that can increase release success is size: released males may benefit competitively from being larger than their field counterparts. However, there could be a risk in releasing only large males if small field females avoid these males and instead prefer small males. Here we investigate this risk by evaluating mating success for mosquitoes differing in size. We measured mating success indirectly by coupling size with *Wolbachia*-infected or uninfected mosquitoes and scoring cytoplasmic incompatibility as a way of estimating relative mating success. Large females showed no evidence of a mating preference, whereas small males were relatively more successful than large males when mating with small females, exhibiting an advantage of around 20-25%. Because field females typically encompass a wide range of sizes while laboratory reared (and released) males typically fall into a narrow size range of large mosquitoes, these patterns can influence the success of release programs which rely on cytoplasmic incompatibility to suppress populations and initiate replacement invasions. Releases could include some small males generated under low food or crowded conditions to counter this issue, although this would need to be weighed against issues associated with costs of producing males of various size classes.

## Background

Aedes aegypti mosquitoes are currently being released around the world for disease suppression. Approaches include replacement strategies aimed at introducing *Wolbachia* infected mosquitoes that directly interfere with viral transmission [1, 2], and population suppression programs that aim to release males that induce sterility through irradiation of males [3, 4] or incompatibility generated through *Wolbachia* [5, 6] which are currently underway (http://www.nea.gov.sg/public-health/environmental-public-health-research/Wolbachia-technology; https://mosquitomate.com/?v=3.0). Other future possibilities include population suppression through the introduction of deleterious endosymbiont effects [7], strategies involving genetically modified mosquitoes [8, 9, 10] or a combination of approaches [11].
In these strategies, it is essential to release mosquitoes that can compete with those in natural populations, facilitating the replacement of one type of mosquito by another and/or the suppression through the induction of male sterility. This can be challenging because released mosquitoes can be at a disadvantage compared to those in natural systems. Various factors including pesticide susceptibility [12], adaptation to favourable laboratory conditions [13, 14], size reflecting nutrition [15, 16], inbreeding [17, 18, 19] and thermal acclimation will influence the ability of released insects to compete with resident populations. So far, most of these effects have not been studied much in the context of mosquito releases, except for body size [20].

The average size of released *Ae. aegypti* mosquitoes tends to be much larger than the average size of those from natural populations, while the variance in size tends to be much smaller [21]. In releases of *Wolbachia* leading to population replacement, released females were 18% larger than those obtained from field collections, with a CV of 8% or more for field mosquitoes compared to <4% for released mosquitoes [21]. This is no doubt a consequence of released mosquitoes being reared under favourable nutrition and temperature conditions. Under these conditions, larvae develop quickly and evenly, ensuring that adult releases involve the largest number of newly-emerged adults possible. When larval densities of *Ae. aegypti* are increased relative to food availability, adult size sharply decreases along with an increased variance in development time [22]. This can in turn slow the rate of *Wolbachia* incursion into populations [23].

Yet while large *Ae. aegypti* males may be at an apparent advantage as they tend to have a greater sperm capacity [24], transfer more sperm to females [16] and have a slower sperm depletion rate [25], there is also the possibility that some degree of assortative mating for size exists in populations as in Drosophila [26, 27, 28]. In other species, there is evidence for mating preference influencing the impact of interventions. Wild populations of *Ceratitis capitata* [29] and *Dacus cucurbitae* [30] altered their mating preference in response to sterile insect releases. Releases of sterile *Culex* tarsalis males also failed to suppress wild populations as the wild and colonized mosquitoes exhibited a preference for their own type [31]. We have therefore explored whether females prefer similar-sized males by taking advantage of cytoplasmic incompatibility as a way of measuring relative mating success (c.f. [32, 33]).

## Methods

### Mosquito strains and colony maintenance

*Ae. aegypti* mosquitoes were reared in an insectary under standard laboratory conditions as described previously [22]. *Ae. aegypti* infected with the *w*Mel strain of *Wolbachia* (w+) were collected from Queensland in 2013, following field releases [1]. Uninfected (w-), wild-type mosquitoes were collected from Queensland in 2016 from areas outside the release zones. To maintain a similar genetic background between strains, wMel-infected females were crossed to uninfected males for at least three consecutive generations before commencing experiments [34].

### Generating large and small mosquitoes

Previous studies have generated small mosquitoes through larval crowding or constant low nutrition [15, 24], however this can greatly increase the variance and duration of larval development [22]. Since we required large numbers and synchronous larval development for the experiments, we altered nutrition during the fourth larval instar to generate mosquitoes of two distinct size classes with similar development times. Mosquitoes for the large size class were provided food ad libitum throughout their development; 500 larvae were reared in trays with 4 L of reverse osmosis (RO) water and provided with >0.5 mg of TetraMin tropical fish food tablets (Tetra, Melle, Germany) per larva per day until pupation. Mosquitoes for the small size class were reared identically to the large size class for the first four days of their development. At this point (at 26°C), most larvae will have committed to pupation but have not yet reached their maximum weight [35, 36]. Therefore, 96 hrs after hatching, larvae for the small size class were transferred to trays with 4 L of fresh RO water and then provided with 0.1 mg of TetraMin per larva per day until pupation. This rearing regime produced adults that developed at approximately the same rate but with two distinct size classes (Figure 1). Pupae were removed from trays daily and placed into round plastic containers with 200 mL of water, and adults from each sex, size class and *Wolbachia* infection type were left to emerge into separate 19.7-L BugDorm-1® cages (MegaView Science Co., Ltd., Taichung City, Xitun District, Taiwan). Adults were matured for at least two days before being used in mating experiments.

**Figure 1.**
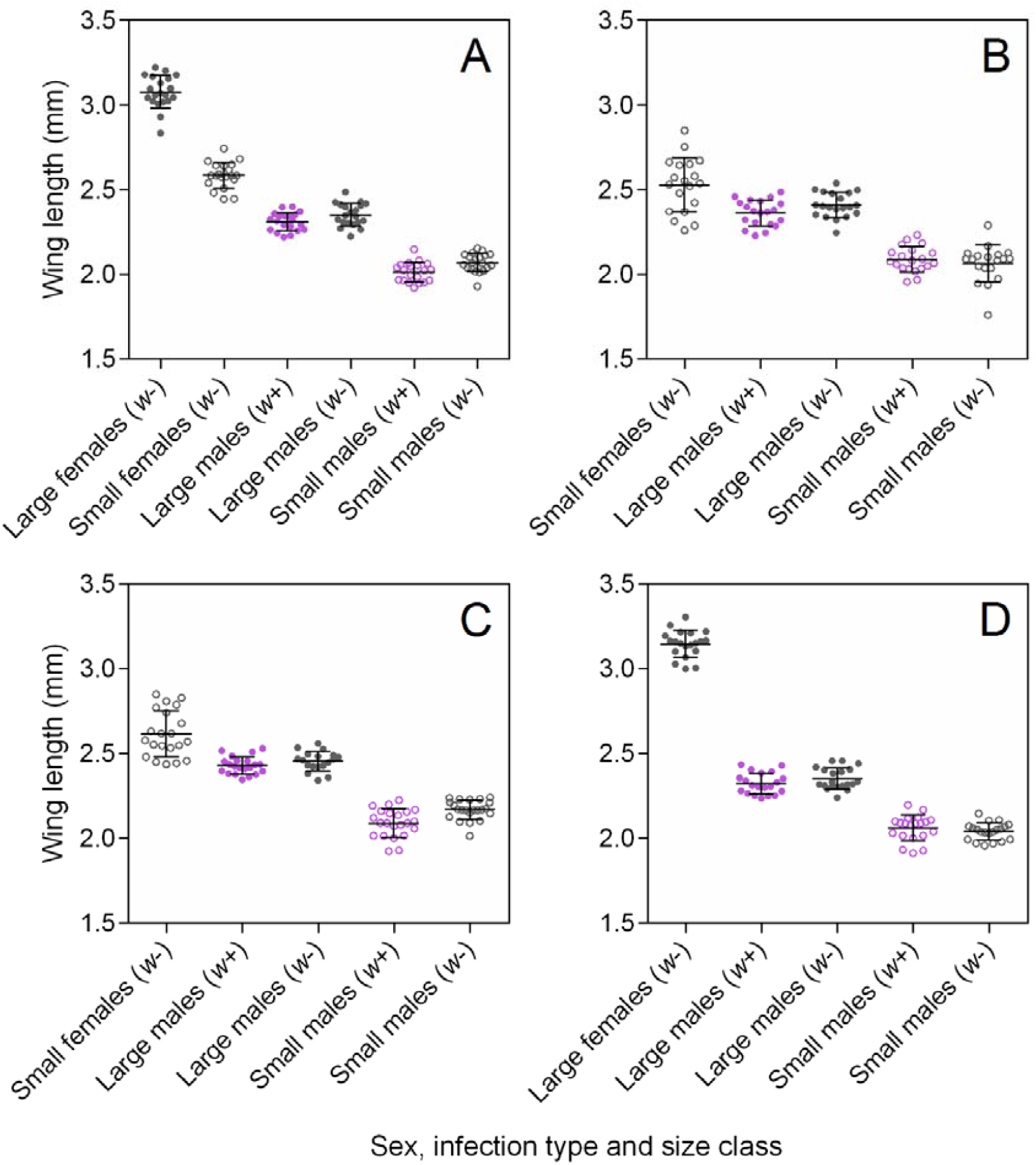
Effect of treatments on wing size in the first (A), second (B), third (C) and fourth (D) experiments. In all cases there was clear separation between the size classes produced by the different rearing conditions, regardless of *Wolbachia* infection type. 20 wings were measured from each group. Error bars are standard deviations.

### Mating

We tested the ability of males from small and large size classes to compete against each other for access to small or large females in laboratory cages. We estimated mating competitiveness by crossing *w*-females with *w*-and *w*+ males held in equal proportions. Under standard laboratory conditions, *w*-females mated to *w*+ males do not produce viable progeny, while crosses with *w*-females and *w*-males produce eggs that are almost all viable [37]. We therefore used egg hatch rate as a proxy for mating success, with higher hatch rates indicating greater competitiveness of the *w*-male [32, 33]. We established eight crosses with *w*-males, *w*+ males and w-females of different size class combinations (Table 1). In each cross, 50 *w*+ and 50 *w*-males were aspirated into a 12 L plastic cage with mesh sides (30 × 20 × 20 cm) and allowed to mix for several minutes before 50 *w*-females were aspirated into the same cage. Five replicate cages were established for each cross.

**Table 1.** List of experimental crosses with w-males, w+ males and w-females of large and small size classes.

| Cross | Female size class | *w*+ male size class | *w*- male size class | Replicates |
| --- | --- | --- | --- | --- |
| 1 | Large | Large | Large | 10 |
| 2 | Large | Small | Small | 10 |
| 3 | Large | Large | Small | 10 |
| 4 | Large | Small | Large | 10 |
| 5 | Small | Large | Large | 15 |
| 6 | Small | Small | Small | 15 |
| 7 | Small | Large | Small | 15 |
| 8 | Small | Small | Large | 15 |

Three days after establishing the cages, females were blood fed, and a single cup filled with larval rearing water and lined with a sandpaper strip was added to each cage. Sandpaper strips were collected daily and photographed, and the number of eggs on each strip was counted in ImageJ using the Cell Counter plugin (https://imagej.nih.gov/ij/plugins/cell-counter.html). Eggs were hatched four days post-collection and larvae were counted four days after hatching. Egg hatch rates were estimated by dividing the total number of larvae by the number of eggs from each cage. Crosses with large females were repeated once, and crosses with small females were repeated two more times in later generations.

### Confirmation of body size and *Wolbachia* infection status

We measured a sample of wings from large and small size classes to confirm that sizes fell into distinct groups. For each experiment, 20 males and 20 females from each size class and infection type were stored in 100% ethanol, and wings were measured according to methods described previously [38]. To confirm the *Wolbachia* infection status of mosquitoes used in the experiments, we screened 30 individuals from each group using a previously described quantitative real-time polymerase chain reaction assay [39].

### Analysis

All data were analysed using SPSS Statistics 24.0 version for Windows (SPSS Inc, Chicago, IL). We ran a linear model with rearing condition and infection as fixed factors to investigate their impact on (untransformed) wing size. Experiment was included as a random factor in this design.

For the hatch rate data, we angular transformed hatch rates and then compared the male size classes. Again, experiment was included as a random factor in the design. We then used the data to compute the relative fitness of the small versus large males based on the proportional changes in hatch rates when large or small males were involved in the matings compared to average hatch rates. For instance, with average hatch rates for a particular class of females of h_ave_, the difference in fitness of small infected males with hatch rates h_s_ relative to all treatments was computed as 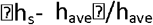, with lower hatch rates indicating an advantage to the smaller males and vice versa.

## Results

We measured the wing length of males and females from all groups to confirm that we generated adults of distinct size classes in each experiment (Figure 1). In the first experiment where we produced females of both size classes, females from the small size class (mean = 2.59 mm, SD = 0.08) were 16% smaller than females from the large size class (mean = 3.08 mm, SD = 0.10, Figure 1A); this difference was highly significant (F_1,38_ = 323.812, P < 0.001). Females from the large size class differed in their wing length between experiments (F_1,38_ = 6.018, P = 0.019), which could indicate differences in rearing conditions across generations, however females from the small size class did not differ across experiments (F_2,57_ = 2.263, P = 0.113).

The different rearing conditions also produced males of two distinct size classes; males from the small size class (mean = 2.08 mm, SD = 0.08) were 12.5% smaller than males from the large size class (mean = 2.37 mm, SD = 0.08, F_1,306_ = 1555.010, P < 0.001). *Wolbachia* infection type had a significant effect on male size (F_1,306_ = 14.311, P < 0.001), though w+ males were only 1.2% smaller than w-males on average. Male wing length also differed across experiments (F_3,306_ = 35.504, P < 0.001), likely reflecting subtle differences in rearing conditions. However, this did not affect our ability to generate adults of two distinct sizes in all four experiments; interactions between rearing condition and experiment were not significant (F_3,306_ = 0.809, P = 0.490).

We compared differences in (arcsine transformed) egg hatch rate between crosses to determine any effects of male and female body size on mating success. For crosses with females from the large size class, egg hatch rate was unaffected by the male size class combination (F_3,32_ = 0.657, P = 0.631), and there was no effect of experiment (F_3,32_ = 4.748, P = 0.117). Thus, mating competitiveness with large females appears to be unaffected by male body size (Figure 2).

**Figure 2.**
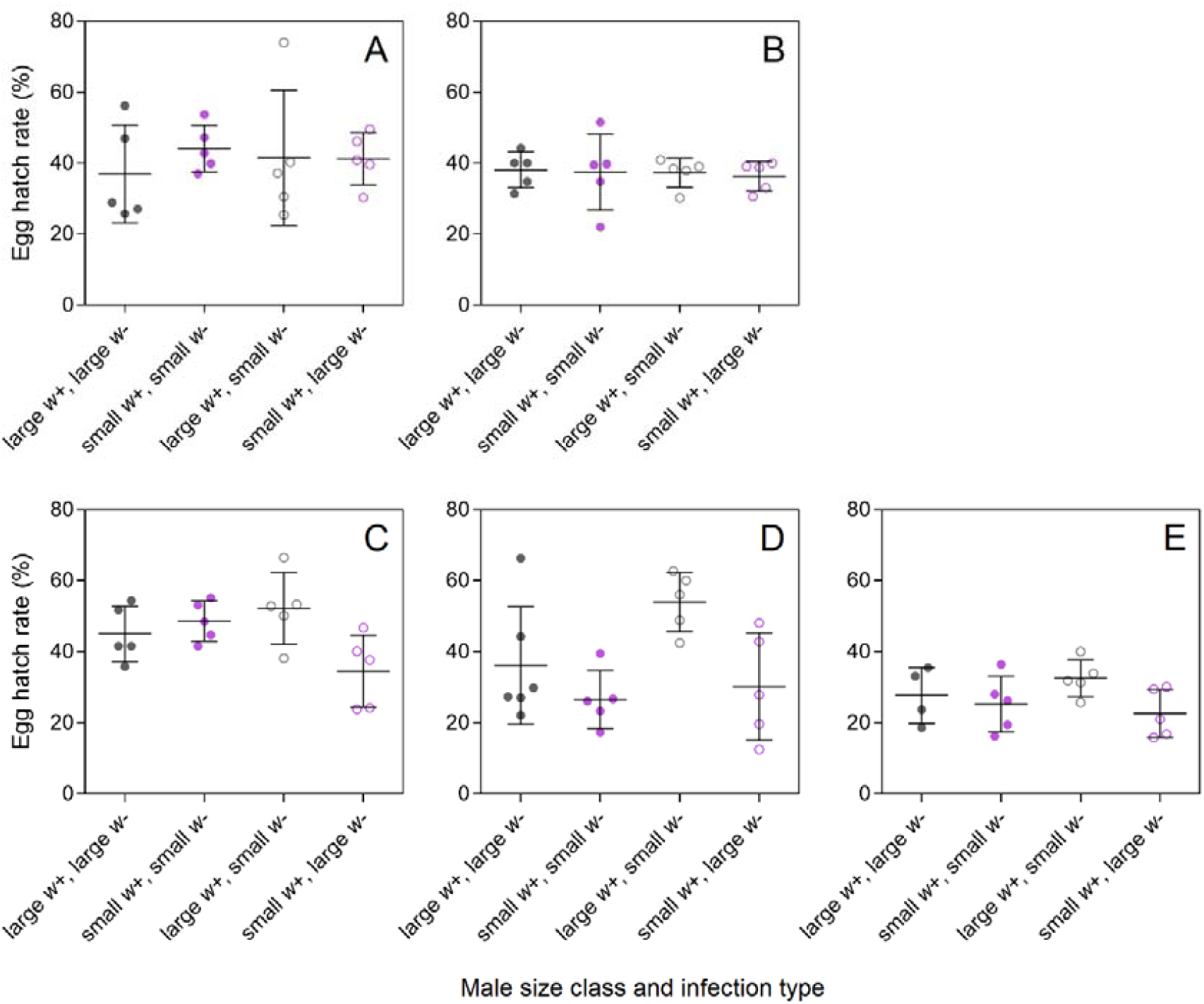
Egg hatch rates from crosses with uninfected females and both infected (*w*+) and uninfected (*w*-) males of different size class combinations. Comparisons were carried out twice for large females (A-B) and three times for small females (C-E). Error bars are standard deviations.

In contrast to large females, we observed a consistent pattern indicating an effect of male body size on mating success with small females (Figure 2). In the first experiment, there was a difference among groups (F_1,16_ = 3.971, P = 0.027) due to lower hatch rates than expected when small infected males competed with large uninfected females (Figure 2). Across all three experiments with small females, there was also a significant effect of male size class combination (F_3,54_ = 7.540, P< 0.001) and there was a significant effect of experiment (F_2,54_ = 14.897, P < 0.001), reflecting the lower hatch rates in the third experiment (Fig. 2). Overall hatch rates can vary between experiments depending on the nature of the eggs laid and drying conditions, but these should be similar for all treatments within an experiment. The small infected males had an advantage over large uninfected males of around 20% (reduced hatch rate relative to overall average) while the small uninfected males had an advantage of around 27%.

## Discussion

Effective mating is key to the success of *Wolbachia* release programs aimed both at population suppression and at replacement. Here we show that where female *Ae. aegypti* are small, there is likely to be an increased mating success of small males. Because females mostly mate only once [40, 41], this results in a mating advantage to small males regardless of whether the males are infected or uninfected by *Wolbachia*. Therefore, although *Wolbachia* have no direct effect on mating [33, 42] except under particularly high or low frequencies [43], environmental effects as used here to generate mosquitoes of different sizes could influence mating success.

Other researchers have pointed to the potential advantages of matching the size of released males to field males [44]. With a mix of males reared under different conditions, it should be possible to approximate the range of sizes typically seen in the field. For instance, by rearing mosquitoes at high densities, Hancock and others [45] produced mosquitoes whose size distribution closely matched what was observed in the field. Given that our experiments were conducted in small cages under laboratory conditions, the extent to which increased mating success of small males will translate to field conditions is unclear. Factors such as dispersal ability could be affected by body size and may influence the ability of males to successfully inseminate females in open field situations. *Large Ae. aegypti* males have distinct advantages relative to small males in other ways such as longevity [20] and sperm capacity [24] that could also contribute to increased mating success under field conditions.

More research on the importance of these factors in field mating success is needed. It would also be interesting to understand the reasons why small females prefer to mate with small males. We suspect that this preference could be due to difficulty in achieving successful mating with large males. In preliminary experiments it appears that more attempts may be required before mating is successful, and mating may tend to be restricted to only some surfaces in a cage (Ross, unpublished). These types of effects could make small females mating with large males more susceptible to predation during mating.

There are likely to be costs associated with modifications to rearing conditions that influence size. When producing males for release, high density/low food rearing usually results in staggered emergence [22], but synchronized emergence is desirable to ensure that males can be efficiently produced. In our experiments we could generate small males with little delay in development time; mass rearing procedures could simply alter the timing of feeding at the fourth larval instar to produce adults of a range of sizes [36]. However, this approach may not be feasible for sterile or incompatible insect programs where only males are released, as sexing pupae by size becomes less reliable. For these programs, it is doubtful whether a (maximum) fitness cost of 20-25% for large males would be countered by extra production costs required to produce small males, unless there was a large density dependent component not measured in this study that would increase the size of such costs.

Nevertheless, there are situations where the current results could be useful in increasing release success. For instance, in replacement strategies, *Aedes aegypti* mosquitoes are often released by placing eggs in containers which contain food and are left outside to produce both males and females. This strategy is currently being used in some *wAlbB* releases in Kuala Lumpur. In this situation where the release containers provide infected mosquitoes over an extended period, food could be limited to ensure that males having a range of sizes emerge from the release containers, however this could also slow the invasion of *Wolbachia* into a population [45].

## Conclusions

We show that small *Ae. aegypti* females exhibit a preference for small males over large males in laboratory mating experiments. Because females in the field typically encompass a wide range of sizes, the release of only large males from the laboratory could affect the success of release programs. Our results are of relevance to modified mosquito release programs, particularly for replacement approaches where sexing by size is not required.

## Supplementary information

Data S1. Raw wing length and hatch rate data presented in figures 1 and 2.

## Declarations

### Ethics approval and consent to participate

Blood feeding of mosquitoes on human subjects was approved by the University of Melbourne Human Ethics Committee (approval #: 0723847). All volunteers provided informed written consent.

### Consent for publication

Not applicable.

### Availability of data and materials

The datasets supporting the conclusions of this article are included within the article and its additional file.

### Authors’ contributions

AGC performed experiments, screened for *Wolbachia* infection and contributed to writing the manuscript. PAR performed experiments, measured wings, analysed data, generated figures and wrote the paper. AAH supervised the experiments, analyzed data and wrote the paper. All authors read and approved the final manuscript.

## Acknowledgements

The authors thank Véronique Paris and Jason Axford for technical assistance.

